# A Far-Red FRET biosensor for AMPK enables multiplexed imaging of single-cell bioenergetic homeostasis

**DOI:** 10.64898/2026.05.13.723899

**Authors:** Nicholaus L. DeCuzzi, Marion Hardy, Markhus B. Cabel, Jason Hu, Christina Abbate, Michael Pargett, John G. Albeck

**Author notes:** These Authors contributed equally (Nicholaus L. DeCuzzi, Marion Hardy).

## Abstract

Metabolic homeostasis has been studied primarily at the tissue and organism level, identifying molecular control mechanisms such as the energy charge-sensing kinase AMPK. Feedback loops involving AMPK and other regulators align cellular ATP generation and consumption, determining energetic balance. Recent work has demonstrated surprising oscillatory dynamics in AMPK activity, revealing unidentified kinetic modulation in single-cell homeostatic behaviour. However, probing the kinetic mechanisms of intracellular feedback requires simultaneous observation of multiple energetic parameters, and such experiments are precluded by the shared wavelength band occupied by most metabolic biosensors. We have overcome this obstacle by constructing a red-shifted FRET-based AMPK activity biosensor, RAMPKAR2, that is comparable to existing FRET-based AMPK activity biosensors. Multiplexed imaging of RAMPKAR2 with PercevalHR, which detects ATP/ADP ratio, confirmed that the kinetics of AMPK activity and ATP/ADP ratio are tightly coupled, with a lag of less than 6 minutes at the single-cell level. Pairing of RAMPKAR with HYlight, which detects the glycolytic intermediate fructose 1,6-bisphosphate (FBP), revealed that glycolytic activity co-oscillates with AMPK, shifted by ∼1.5 hours, and that these oscillations are suppressed by sustained AMPK activity. Together these data advance a model in which temporally offset increases in glycolytic ATP supply and AMPK deactivation contribute to single-cell oscillations.

## Main text

As the pool of excess ATP provides the primary “energy charge” or driving potential for biochemical reactions, its regeneration from ADP is essential in all cells. Furthermore, the stability of the driving potential relies on the balance between the production of ATP and its consumption at any given time via a regulatory system that remains understood only in part. ATP generation is carried out primarily by glycolysis and oxidative phosphorylation (OxPhos). Cells regulate these ATP-generating pathways alongside ATP-depleting pathways to align ATP production with the availability of nutrients and cellular energy demands^1,2^. This coordination is known to depend in part on a key signaling protein, AMP-activated protein kinase (AMPK), which is activated by decreased energy charge (i.e. ratio of ATP to ADP and AMP)^3,4^. AMPK is a heterotrimeric protein kinase; its regulatory gamma subunit binds ATP, ADP, and AMP, enabling it to provide a readout of cellular energy charge, which is fundamentally sensitive to changes in glycolysis or OxPhos^5^.

Recent studies have shown that glycolysis, OxPhos, and AMPK activity can vary substantially on the single-cell level, in both living tissues and cell culture models^6–8^. For example, oligomycin, a specific inhibitor of the F0/F1 ATPase complex that blocks the generation of ATP by mitochondria, induces immediate AMPK activation, which is then followed by adaptive behaviours that play out over many hours, including repeated oscillations^6,7,9^. These extended dynamic changes reflect the operation of feedback loops that restore cellular energy charge (indicated by decreasing AMPK activity). Dynamic AMPK activation is also observed under baseline conditions, indicating that such feedback loops operate continually in response to cell-intrinsic needs^10^. It remains unclear which feedback loops are responsible for these observed kinetics, although there are numerous candidates, such as the allosteric modulation of phosphofructokinase-1 by AMP and ATP^11^, or the suppression of mTOR-driven protein synthesis by AMPK-mediated phosphorylation^12^. Cell-to-cell heterogeneity observed in these studies reveals that there are preexisting subpopulations with different metabolic configurations, and that the process of resolving metabolic limitations can take different dynamic paths. By studying such kinetics in depth, it may eventually be possible to develop quantitative models of metabolic self-regulation that explain the heterogeneity of cells and predict the metabolic behaviour of tissues^13,14^.

Single-cell studies rely on genetically encoded fluorescent biosensors, which may be imaged in live cells to report the abundance of key metabolites or the activity of metabolic regulatory proteins. Biosensors are currently available for intracellular glucose^15,16^, ATP:ADP ratio^17^, NADH:NAD+ ratio^10,18^, glycolytic intermediates^19–21^, related metabolites^21,22^, and the regulatory activity of AMPK^6,9,23–25^. Because the kinetics of metabolite concentrations and AMPK activity vary considerably between cells, performing simultaneous measurements of these variables within a single cell is critical to establish the temporal offsets that govern homeostatic feedback mechanisms. At present, a key barrier to such experiments is the spectral overlap of the vast majority of available fluorescent biosensors, which employ either a CFP/YFP FRET pair or a circularly permuted GFP (cpGFP) and occupy the same region of the visible spectrum^10,17,18,21,24,26^. Although spatial and temporal multiplexing strategies are available^27,28^, spectrally compatible sensors would provide a more straightforward, easily adopted, and broadly practical toolset.

We addressed this barrier by developing a red-shifted FRET AMPK biosensor, based on the existing AMPKAR2^7^, that is spectrally compatible with both CFP/YFP FRET and cpGFP-based biosensors (**Figure 1A**). Red AMPKAR2 (RAMPKAR2) was constructed following a previous strategy^29^, by replacing the mTurquoise2 (cyan) and mYPet (yellow) fluorophores of the existing FRET-based AMPK biosensor (AMPKAR2)^7^ with the far red fluorescent proteins miRFP720^30^ and miRFP670nano3^31^, respectively. To compare the activity of the red-shifted sensor with the original, we generated MCF10A cells expressing both RAMPKAR2 and AMPKAR2 (**Figure 1B**). Live imaging was performed for both biosensors concurrently, during which we treated cells with either the AMPK activator MK8722^32–35^ or a vehicle control (**Figure 1C**). To quantify the response to treatment, the fraction of reporters in an active configuration (*E*f_A_) was computed via measurement of the ratiometric signals for each biosensor signal per cell and time point. Differences in biosensor signals before and after treatment (ΔAMPKAR2 or ΔRAMPKAR2) were then computed for each cell (**Figure 1D**). Vehicle treatment caused no significant change (p-value > 0.05) for either AMPKAR2 or RAMPKAR2 (δ of -0.008 and 0.003 *E*f_A_, respectively), while MK8722 significantly (p-value = 0.0018) increased both signals (signal deltas of 0.03 and 0.05 *E*f_A_, respectively), demonstrating very similar responses to AMPK activation. While AMPKAR2 appeared to be sensitive to vehicle spikes, RAMPKAR2 did not display a similar sensitivity, which could be due to a variety of factors, including fluorophore sensitivity to changes in the pH of the imaging medium caused by the treatment. While RAMPKAR2 had the stronger response, it is also the noisier of the signals, observed via greater signal variance in over time (**Figure S1A**), suggesting there remains room for optimization of its readout properties.

**Figure 1:**
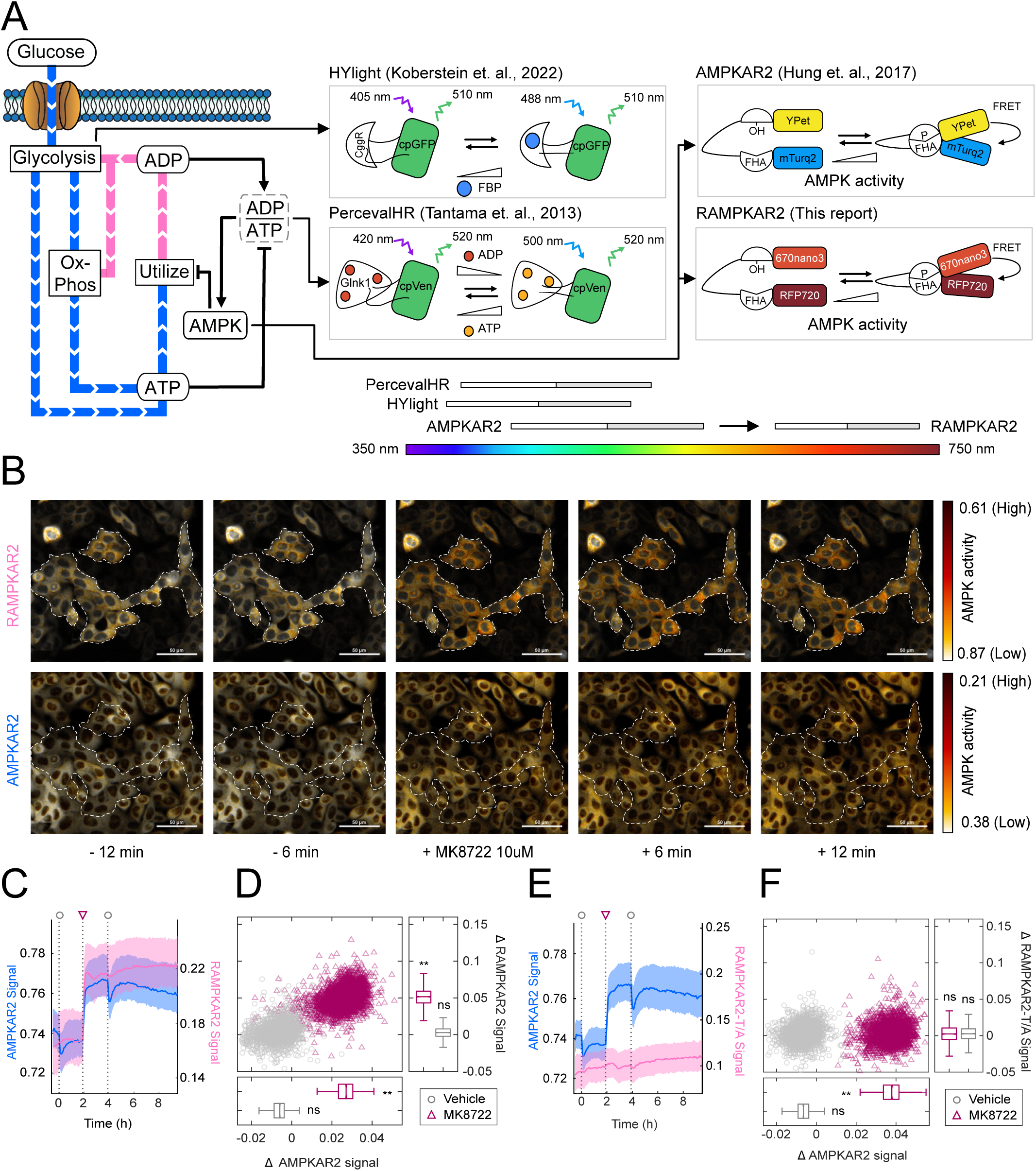
Validation of the Red-FRET AMPK biosensor, RAMPKAR2. **(A)** Schematic diagram of metabolic processes and related biosensors. RAMPKAR2 is a modified AMPKAR2 in which YFP (Ypet) and CFP (mTurq2) were, respectively, replaced with miRFP670nano3 (670nano3) and miRFP720 (RFP720), shifting its excitation and emission properties into the red/far-red spectrum. **(B)** Example time-lapse images of MCF-10A cells co-expressing RAMPKAR2 and AMPKAR2, before and after treatment with the AMPK activator MK-8722. Paired images of the RAMPKAR2 FRET:donor ratio (top row) or AMPKAR2 CFP:YFP ratio (bottom row) are coloured according to the scales at right. The same field of cells is shown in both top and bottom rows, with the outlined region indicating groups of cells expressing both biosensors. Scale bar: 50 μm. **(C)** Mean AMPKAR2 and RAMPKAR2 signals (blue/pink lines) with 25^th^ to 75^th^ interquartile range (IQR, shaded light blue/pink regions) from MCF-10A cells co-expressing the 2 biosensors. Plots shown are representative data from 1 of 3 experimental replicates, each having >250 cells per condition. Coloured shapes indicate treatments applied to the cells as indicated in legend. **(D)** Scatter plots of single cell changes (Δ) in AMPKAR2 and RAMPKAR2 signals from 1 hour before treatment to 1 hour after treatment with either vehicle (grey, circles) or MK8722 (purple, triangles) treatment. Accompanying box and whisker plots show medians and IQRs of scatter plot data for each respective biosensor and treatment. Significance of responses was assessed by one-sample t-test of the mean delta value against zero, across experimental replicates (n=3 independent experiments, each comprising 4 technical replicates with >800 cells per condition). (ns = not significant, ** = p-value <0.002). **(E,F)** Comparison of the phosphoacceptor mutant RAMPKAR2-T/A to AMPKAR2, plotted as in (C) and (D), respectively.

We verified that RAMPKAR2 did not acquire any unintended off-target phosphorylation sites by generating RAMPKAR2-T/A, which contains a threonine-to-alanine substitution in the AMPK substrate recognition domain. This T/A mutation eliminates the AMPK phosphorylation target within the biosensor and, as expected, the mutant does not exhibit a change in FRET signal even when cells bearing it are treated with MK8722 (**Figure 1E-F, timelapse images in Sup Figure 1A**). Conversely, the co-expressed AMPKAR2 maintained its responsiveness in these same cells (signal delta -0.007 vs. 0.037 *E*f_A_, for vehicle vs. MK8722).

In previous studies, we found that cellular responses to oligomycin are highly heterogeneous, with both ATP:ADP ratio (per the ATP:ADP ratiometric biosensor, PercevalHR) and AMPK activity (per AMPKAR2) displaying individual cell behaviours ranging from no apparent change to a stable change in the signal, to oscillations with a roughly 3-hour period^6,7^. However, due to the issue of spectral compatibility, we were unable to observe the response of AMPK to the change in ATP:ADP ratio in an individual cell. We therefore generated MCF10A cells expressing both RAMPKAR2 and PercevalHR^7,17^. At the population average level, following oligomycin treatment, we observed an immediate decrease in PercevalHR signal concomitant with a peak in RAMPKAR2 (**Figure 2A, S2A**), while vehicle treatment induced only small transient peaks in both reporter signals. At reduced glucose concentrations, the magnitude and duration of the decrease in ATP:ADP signal was greater, while AMPK activation was correspondingly more prolonged. This correspondence is consistent with our expectation that at lower glucose concentrations, glycolysis is less effective in compensating for the inhibited ATP production from OxPhos, resulting in a longer period of reduced energy charge.

**Figure 2:**
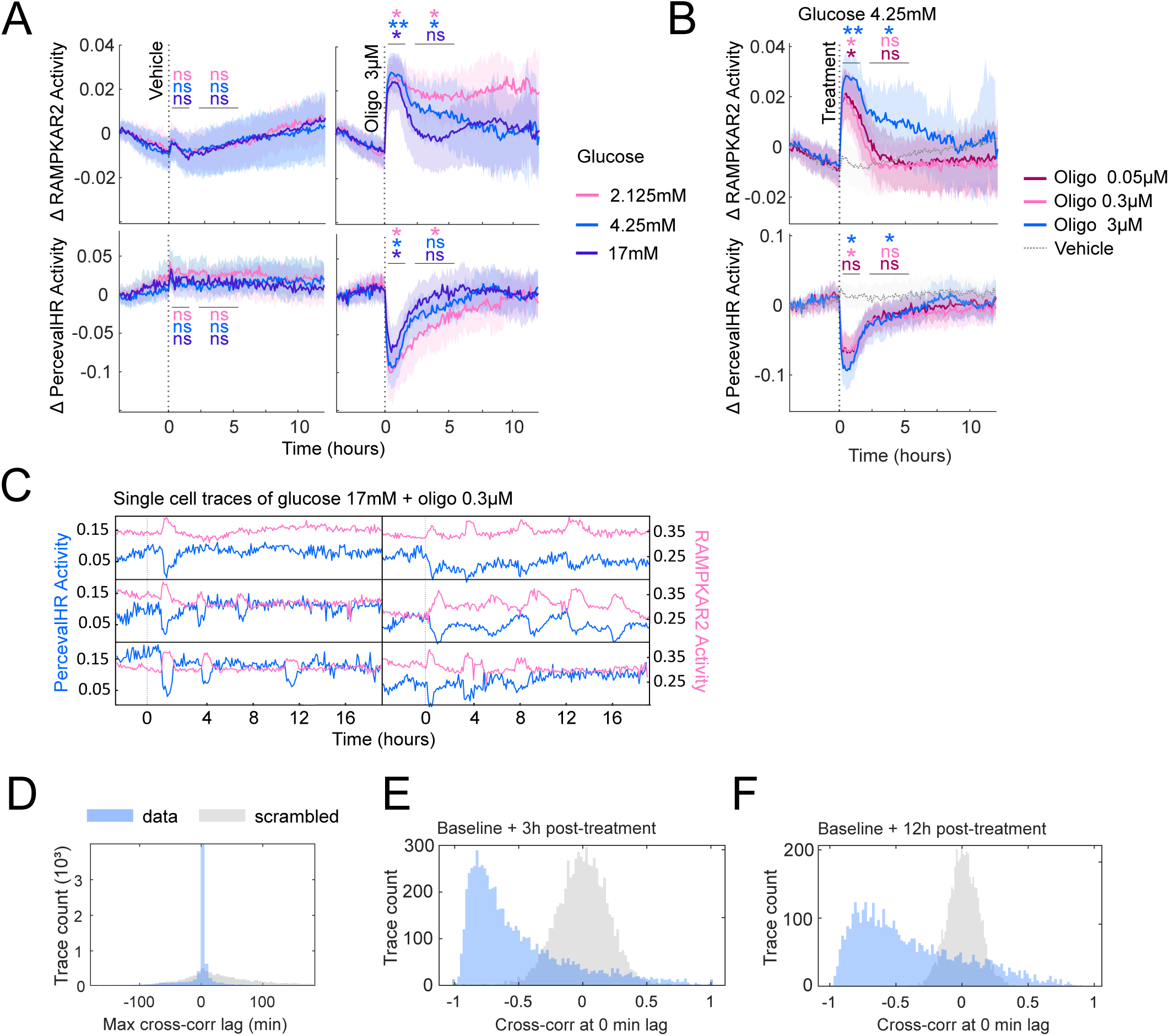
Multiplexed measurements of AMPK activity and ATP:ADP ratio. **(A)** Mean Δ RAMPKAR2 (top) and Δ PercevalHR (bottom) signal in response to 3 μM oligomycin under varying glucose concentrations. MCF-10A cells were first cultured in 17, 4.25, or 2.125 mM glucose and treated with 3 μM oligomycin at the indicated time. Shaded regions show 25^th^ to 75^th^ IQR of individual cell measurements. Sensor traces were baseline-normalized to the first time point and are shown as Δ sensor activity. PercevalHR signal is shown as ATP:ADP ratio (GFP/Sapphire). **(B)** Mean Δ RAMPKAR2 (top) and Δ PercevalHR (bottom) responses to oligomycin in 4.25 mM glucose. PercevalHR-RAMPKAR2 MCF-10A cells were first cultured in 4.25 mM glucose for 8 hours and treated with 0.05, 0.3 or 3 μM oligomycin at the indicated time. Shaded regions show IQR. Signals are baseline normalized and are shown as Δ sensor activity. For both A and B, paired t-tests comparing the mean RAMPKAR2 activity within the indicated time ranges were performed across experimental replicates (n = 3) where ns (non-significant), * = p-value < 0.05, and ** = p-value < 0.005 (with colour respective to the condition). All vehicle controls were non-significant. **(C)** Six example single cell tracks from PercevalHR-RAMPKAR2-expressing MCF-10A cells in 17 mM glucose, treated with 0.3 μM oligomycin. **(D)** Maximum cross-correlation lag in minutes between the PercevalHR and RAMPKAR2 signals per single cell trace (blue bins). Data were scrambled to generate a control distribution (grey bins). **(E)** Cross-correlation values at a fixed 0-minute lag between the PercevalHR and RAMPKAR2 traces spanning 4 hours before and 3 hours after oligomycin treatment, across glucose concentrations. **(F)** Cross-correlation values at a fixed 0-minute lag between the PercevalHR and RAMPKAR2 traces spanning 4 hours before and 12 hours after oligomycin treatment, across glucose concentrations. For A-F, data were pooled from 3 experimental replicates (n = 3), totaling >6,500 cell traces.

When oligomycin concentration was varied from 0.05 to 3 μM, we saw little difference in the ATP:ADP ratio response (**Figure 2B**). However, the duration of the AMPK activity response increased across these conditions from 2 hours to >5 hours, indicating more persistent impairment of energetic balance, on average. At the single-cell level, we observed heterogeneity in activity over time where cells displayed 3 major subtypes of kinetics. In one subset of cells, ATP:ADP ratio and AMPK activity measurements displayed opposing, apparently phase-locked oscillatory behaviour with a period of 3-4 hours (denoted S1 in **Figure 2C**). A second subpopulation (S2) displayed a single peak in AMPK activity and a corresponding single depression in ATP:ADP ratio, with both signals resolving to baseline levels at similar rates. Finally, a third subset of cells (S3) displayed multiple peaks but with less clearly defined oscillations. We evaluated the overall relationship between PercevalHR and RAMPKAR2 signals using cross-correlation of the paired biosensor signals from individual cells. This analysis revealed a narrow distribution of lag times centered at 0 (**Figure 2D**). With the lag fixed at 0, correlation values during the 3 hours following treatment were distributed heavily toward negative values, with a median of -0.64 and a mode of -0.83 (**Figure 2E**), indicating strongly opposing time series behaviour. Across a larger time window, a strong tendency toward negative correlation persisted, but was less heavily skewed (-0.47 median; -0.79 mode) (**Figure 2F**). This analysis confirms at the single-cell level that AMPK activation is essentially simultaneous with decreases in ATP/ADP ratio, to the extent observable with the sampling frequency of our experiment (6 minutes), consistent with AMPK’s direct regulation by allosteric binding to adenosine nucleotide pools. The increasing spread of correlation values over longer time windows indicates some combination of (1) the signals diminishing compared with measurement noise and (2) potential contribution from other regulatory inputs to AMPK.

One mechanism for AMPK to restore energetic homeostasis is by accelerating glycolysis^36–38^, a connection also implicated in previous analyses that observed heterogeneity in AMPK activity^7^. We further investigated this idea by generating cells co-expressing RAMPKAR2 and HYlight, a cpGFP-based sensor for the glycolytic intermediate fructose 1,6-bisphosphate (FBP)^19^. As before, these cells were challenged with oligomycin during live imaging, but in contrast to Perceval, the average HYlight signal showed a brief decrease lasting 20-30 minutes, followed by a steady increase over 1-6 hours and an eventual plateau phase (**Figure 3A**). In evaluating the HYlight signal, it is important to note that a rise in FBP is *not* synonymous with an increase in glycolysis, but rather with a shift in the relative flux of reactions affecting FBP. The rate of increase in HYlight signal was linked to the extracellular glucose concentration, likely because with higher glucose availability, greater increases in glycolytic flux can be achieved to compensate for the loss of ATP production from OxPhos. The brief initial decrease in HYlight was most pronounced at lower glucose concentrations and may reflect a transient depletion of FBP as flux in glycolytic reactions accelerates, creating a period in which glucose uptake to supply FBP lags behind FBP usage. This interpretation is further supported by the observation that the HYlight signal remained lower for a longer time in cells treated with higher concentrations of oligomycin (**Figure 3B**).

**Figure 3:**
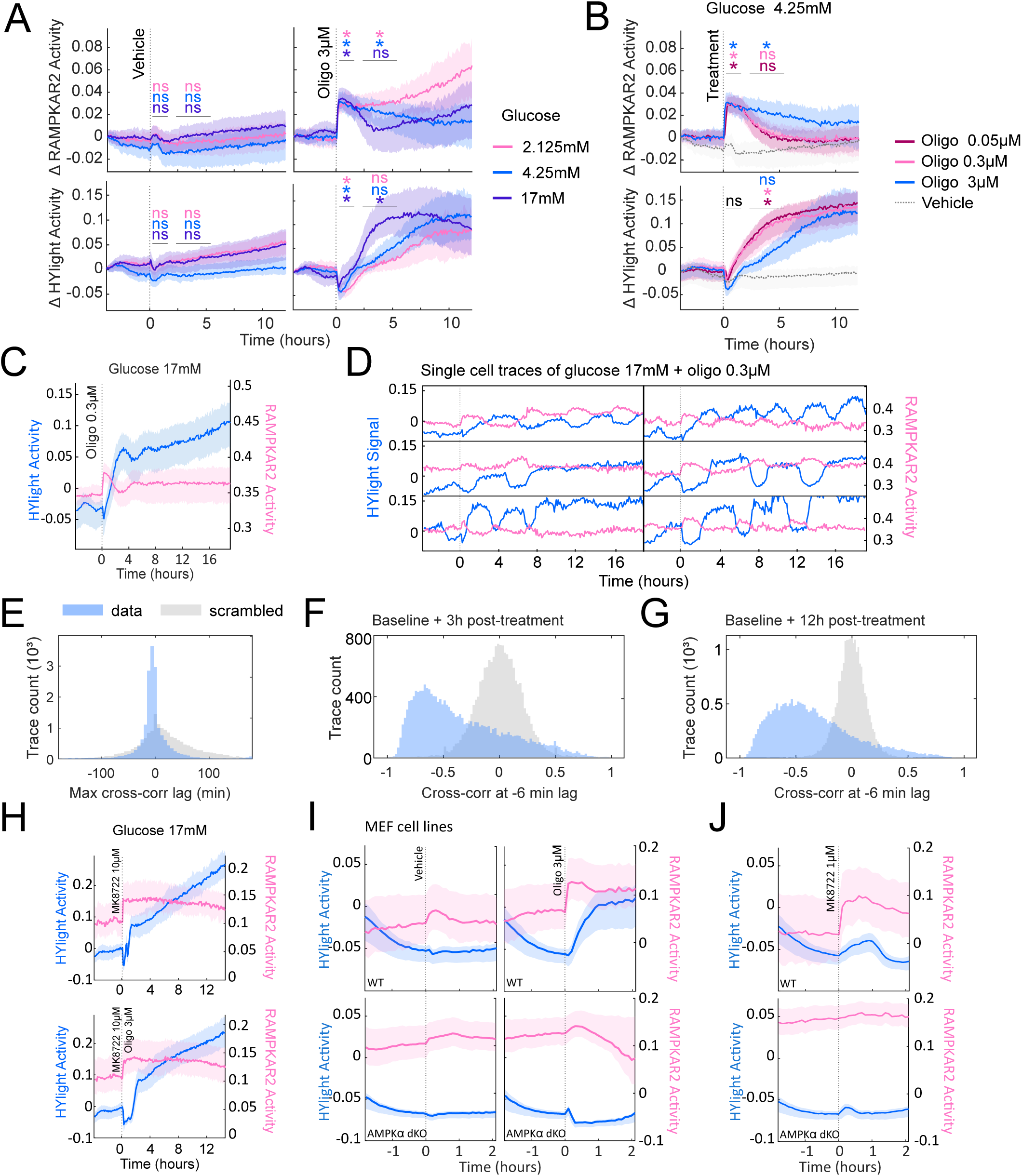
Multiplexed measurements of AMPK activity and FBP levels. **(A)** Mean ΔRAMPKAR2 (top) and ΔHYlight (bottom) signals in 3 μM oligomycin and different glucose concentrations. MCF-10A cells were first cultured in 17, 4.25, or 2.1 mM glucose and treated with 3 μM oligomycin at the indicated time. Shaded regions show 25^th^ to 75^th^ IQR of individual cell measurements. Sensor traces were baseline-normalized to the first time point and are shown as Δ sensor activity. **(B)** Mean ΔRAMPKAR2 (top) and ΔHYlight (bottom) response to oligomycin in 4.25 mM glucose. MCF-10A cells were first cultured in 4.25 mM glucose for 8 hours and treated with 0.05, 0.3 or 3 μM oligomycin at the indicated time. Shaded regions show 25^th^-75^th^ IQR of individual cell measurements. Signals are baseline-normalized and shown as Δ sensor activity. For both A and B, paired t-tests comparing mean RAMPKAR2 or HYlight signals in the indicated time ranges (horizontal bars) to the mean baseline signal were performed across experimental replicates (n = 3), where ns indicates non-significant and * (with the colour respective to the condition) indicates p-value < 0.05. In (B), all vehicle controls were non-significant. **(C)** Average RAMPKAR2 (pink) and HYlight (blue) signal in MCF-10A cultured in 17 mM glucose and treated with 0.3 μM oligomycin. Shaded regions show 25^th^-75^th^ IQR of individual cell measurements. **(D)** Six example single-cell tracks from HYlight (blue) and RAMPKAR2 (pink)-expressing MCF-10A cells in 17 mM glucose and treated with 0.3 μM oligomycin. **(E)** Maximum cross-correlation lag in minutes between the HYlight and RAMPKAR2 signals per single cell trace (blue bins). Data was scrambled as a control to generate a null distribution (grey bins). **(F)** Cross-correlation values at a fixed -6 minute lag between the HYlight and RAMPKAR2 traces, spanning the 4 hours before and 3 hours after oligomycin treatment, across glucose concentrations. **(G)** Cross-correlation values at a fixed -6 minute lag between the HYlight and RAMPKAR2 traces spanning the 4 hours before and 12 hours after oligomycin treatment, across glucose concentrations. **(H)** Average RAMPKAR2 (pink) and HYlight (blue) activity in MCF-10A cultured in glucose 17 mM and treated with MK8722 10 μM and/or oligomycin 3 μM. Shaded regions show 25^th^–75^th^ IQR. **(I, J)** Average RAMPKAR2 (pink) and HYlight (blue) activity in MEF WT (top) or AMPKα1α2 double knockouts (bottom) cultured in 17 mM glucose and treated with vehicle or 3 μM oligomycin (I) or 1 μM MK8722 (J). Shaded regions show 25^th^–75^th^ IQR. Data in (A-H) were collected from MCF-10A cells bearing both the RAMPKAR2 and HYlight sensors. Experiments include 5 experimental replicates (n = 5), >17,000 cell traces. Figures were plotted from representative replicates. Data in (I-J) were collected from MEF cells bearing both the RAMPKAR2 and HYlight sensors. Experiments include 2 experimental replicates (n = 2) with > 5400 cell traces. Figures were plotted from one representative experimental replicate.

An analysis of individual cells further supported our interpretation of the mean AMPK and FBP signals. We first noted that oscillatory behaviour was particularly prominent, and visible even at the population level, in cells treated with 17 mM glucose and 0.05-0.3 μM oligomycin (**Figure 3C-D**). Within these conditions, AMPK activity varied from cell to cell but trended toward oscillatory behaviour. Qualitatively, peaks in AMPK activity appear matched by decreases in HYlight within individual cells, while periods of stable AMPK activity were also stable in HYlight. Cross-correlation analysis estimated that the anti-correlated HYlight signal lags AMPK by roughly 6 minutes on average (21% of cells, with 17.1% and 16.5% lagging 0 and 12 minutes respectively; **Figure 3E**). At a fixed lag of 6 minutes, single-cell correlation values were distributed with a mode of -0.45 and a median of -0.65, softening to a mode of -0.40 and median of -0.45 when considering signals 12 hours after treatment (**Figure 3F-G**).

To gain increased resolution into the relationship between AMPK activation and FBP changes, we treated RAMPKAR2/HYlight MCF-10A cells with the activator MK8722, which produced an immediate increase in RAMPKAR2 signal to a sustained high level, as expected (**Figure 3H, S3B**). The HYlight signal, however, initially dropped for ∼1 hour, sharply increased, then took on a slow and sustained increase that continued for the remaining 12 hours of the experiment. Under co-treatment with MK8722 and oligomycin, the reporter kinetics were similar, but the delay between the initial drop in HYlight and the sharp increase was extended to ∼2 hours. In light of the ∼6 min anti-correlated lag we have observed (**Figure 3E,F,G**), this delay between a step response in RAMPKAR2 and in HYlight suggests that the AMPK and HYlight signals are fundamentally positively correlated but with a lag of approximately a half cycle (i.e. approximately 1.5 hours plus 6 minutes). Such a lag would be consistent with an AMPK-induced increase in glucose uptake and processing through slower mechanisms, such as gene expression^39^.

To test the causal role of AMPK in the FBP shifts indicated by HYlight, we generated two mouse embryonic fibroblast (MEF) biosensor cell lines with RAMPKAR2 and HYlight, based on either wild-type control MEFs (WT-RH) or MEFs lacking both AMPK kinase domain subunits^40^ (α1/α2-KO-RH). We verified that α1/α2-KO-RH cells do not have detectable levels of AMPK activity using immunofluorescence for pACC, a canonical AMPK target, following high-energy stress conditions (**Figure S2**). Under oligomycin treatment, WT-RH MEFs showed an immediate increase in RAMPKAR2 signal and, following a 10-minute delay, an increase in HYlight (**Figure 3I**). In contrast, oligomycin-treated α1/α2-KO-RH MEFs showed a very muted increase in RAMPKAR2 signal (similar in magnitude to vehicle treatment), and after a 10-minute delay, a decrease in HYlight. In WT MEFs, MK8722 induced a slow, moderate, increase in HYlight signal, which lasted approximately 80 minutes and was largely attenuated in α1/α2-KO MEFs (**Figure 3J**). Overall, these data are consistent with the role of AMPK in accelerating glycolysis, albeit with markedly less delay and slower kinetics than observed in MCF10As, which indicate that the regulatory systems in the two cell lines are not poised identically. In particular, the immediate decrease of FBP in MCF10As treated with MK8722 is surprising from a conceptual perspective, as direct phosphorylation of PFKFB3 by AMPK would be expected increase the production of fructose 2,6-bisphosphate, which would then drive increased FBP production through allosteric activation of PFK. As with oligomycin treatment, the transient shift downward could reflect imbalances in the rates of glycolytic reactions before a sustained increase in FBP is established; however, it is unclear in this case if such an imbalance is driven by unrecognized activities of AMPK or of MK8722 itself. Regardless, we interpret these data as supporting the consensus model in which changes in energy charge triggered by OxPhos inhibition drive a nearly immediate response in AMPK activity, which then drives a change in glycolytic flux over a longer period of time through both transcriptional and post-translational mechanisms^4^.

Finally, we asked whether the heterogeneity in single-cell behaviours could provide insight into the mechanisms of oscillatory AMPK dynamics. We first quantified cells based on the number of peaks observed in HYlight activity over the course of the experiment. Under vehicle conditions, cells with 0, 1, or 2 peaks were most common, but we also observed ∼2% of cells with 3 or more peaks, consistent with previous observations of metabolic fluctuations even under basal conditions^7^. Treatment with oligomycin led to an increase in cells with 3+ peaks, up to >25% of the population (**Figure 4A**). As previously observed, oscillating cells were more common at higher levels of glucose^7^ (**Figure 4B**).

**Figure 4:**
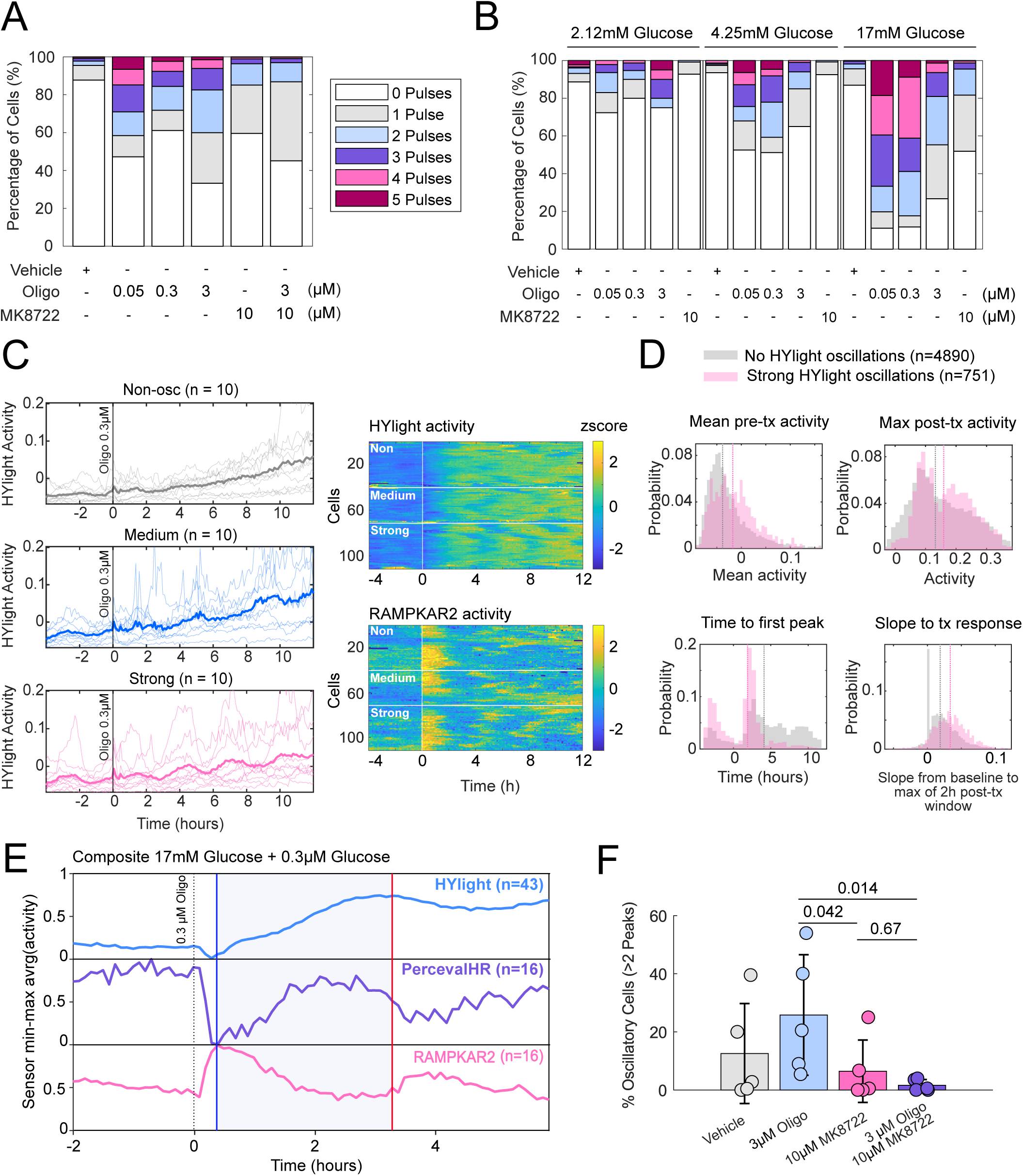
Characterization of HYlight oscillations. **(A)** Stacked percentages of HYlight pulses per treatment. Pulses were quantified from RAMPKAR2-HYlight MCF-10A cells across a 16-hour window (4 hours pre-treatment to 12 hours post-treatment), and data were pooled across glucose concentrations from five experimental replicates (n = 5). **(B)** Stacked percentages of HYlight pulses per glucose and per treatment. Pulses were quantified from RAMPKAR2-HYlight MCF-10A cells across a 16-hour window (4h pre-treatment to 12h post-treatment) and data were pooled from three experimental replicates (n = 3). **(C)** Representative single-cell HYlight traces classified as non-oscillatory, medium, or strong oscillators (left panels), with 10 randomly selected traces per category shown for MCF-10A cultured in 17 mM glucose and treated with 0.3 µM oligomycin. Spatial heatmaps of z-scored HYlight and RAMPKAR2 activity (right panels), one row per cell, ordered by trace similarity. The top 40, middle 40, and bottom 40 rows correspond to non-oscillatory, medium, and strong oscillators respectively. Warm colours indicate higher relative activity. Data was selected from one representative of three experimental replicates. **(D)** Comparative probability histogram of HYlight timeseries features between non-oscillating cells (grey, 4890 cells) and strong oscillators (pink, 751 strong oscillators). Data were pooled across glucose concentrations and oligo treatments (tx) from three experimental replicates (n = 3). **(E)** Composite plot of RAMPKAR2-Perceval and HYlight traces from cells classified as strong oscillatory for either Perceval (16 traces) or HYlight (43 traces) in MCF-10A cultured in 17 mM and treated with 0.3 μM oligomycin. Average population signal was normalized by linear re-scaling (min = 0, max = 1) to make traces comparable. Shown data is from one of three representative experimental replicate. **(F)** Bar plot showing the mean percentage of oscillatory cells per treatment condition. Each bar represents the mean across experimental replicates (n=5), with individual replicate values overlaid as dots. Error bars indicate ±1 SD. Statistical comparisons were performed using a Kruskal-Wallis test on replicate-level proportions with Bonferroni correction for pairwise post-hoc comparisons.

Subpopulations of cells were distinguished by developing a machine learning-based classifier for oscillatory signals, which we used to compare biosensor metrics between highly oscillatory and other cells (**Figure 4C)**. FBP levels in highly oscillatory cells exhibited, on average, higher peak values and steeper rises following oligomycin treatment (**Figure 4D**), suggesting that higher glycolytic rates are more supportive of oscillatory behaviour. Using data from the annotated oscillatory cells of both reporter configurations (RAMPKAR2/Perceval and RAMPKAR2/HYlight), we assembled a composite estimate of the dynamics of AMPK activity, ADP/ATP ratio, and FBP together (**Figure 4E**).

These multiplexed data add new details to our previous conceptual model of oligomycin-induced oscillations^6,7^. After oligomycin treatment triggers an increase in glycolysis, stimulated in part by rising AMPK activity, AMPK shuts off as energy charge rises, due to the tight coupling of AMPK to energy charge (**Figure 4E**, blue line). Although HYlight continues to rise to reach its peak at about 3 hours, consistent with delayed kinetics relative to AMPK that we observed in **Figure 3F**, we suggest that during this time (blue zone) the reduced AMPK activity allows ATP-consuming processes such as protein synthesis to resume, creating additional energetic demand, as we previously showed that protein synthesis has a significant impact on the energetic state of oligomycin-treated cells^6^. We hypothesize that in this low-AMPK state, even an enhanced rate of glycolysis becomes increasingly unable to maintain cellular energy charge, initiating the next cycle and causing AMPK to rise again (**Figure 4E**, red line). This model would explain why a higher glycolytic rate is associated with oscillatory tendency (**Figure 4B**), as higher glycolysis would lead to an earlier inactivation of AMPK (and thus more time for acceleration of energy consumption in the blue zone). Further supporting the model, co-treatment of cells with oligomycin and MK8722, which sustains high AMPK activity, strongly suppresses oligomycin-induced oscillations (**Figure 4F**).

In summary, we report the construction of a new biosensor, RAMPKAR, that enables unprecedented insights into the kinetics of cellular homeostasis through pairing with other metabolic biosensors. This tool allows us to develop new models for cellular bioenergetics that are significant in establishing the identity and timing of processes operating to maintain cellular ATP homeostasis under conditions of mitochondrial failure. Impairment of mitochondrial function is a common thread in many pathological conditions, including aging, neurodegenerative diseases, and exposure to toxins and there is a key need for therapies that ameliorate such damage. In particular, the oscillatory nature of this adaptation is critical, as it implies that the timing of interventions is an important factor in supporting the metabolic functions of damaged cells.

While the model fits our current observations, there are key aspects that remain to be fully elucidated. Perhaps most importantly, while HYlight provides a signal related to glycolytic flux, we cannot rely on it as a direct indication of the rate of ATP production by glycolysis. It will also be important to learn through which mechanisms AMPK induces a sustained increase in FBP, and why FBP reaches a plateau and then falls around 3 hours. Simultaneous imaging with additional sensors, such as those for lactate^20^, pyruvate^21^, NADH^10,18^, or protein synthesis^41^ could provide additional insight, but these remain technically challenging due to their spectral overlap. Integrating these data with computational models of glycolysis^14,42,43^ will provide quantitative insights into bioenergetic regulation. RAMPKAR2 offers a key step toward tracking multiple metabolic processes in a single cell, but additional innovations in sensor design and multiplexing will be needed to fully establish the mechanisms driving oscillatory dynamics.

## Methods

### Key Resources Table

| REAGENT or RESOURCE | SOURCE | Item # |
| --- | --- | --- |
| **Chemicals, Peptides, and Recombinant Proteins** |  |  |
| Oligomycin A | Sigma | 75351 |
| MK8722 | MedChemExpress | HY-111363 |
| Collagen I, Rat tail | Gibco | A1048301 |
| FuGene4K | Promega | E5911 |
| **Experimental Models: Cell Lines** |  |  |
| MCF10A-5e | Janes et al., 2010 | CVCL_0598 |
| MCF10A-AMPKAR2 | Hung et al., 2017 | N/A |
| MCF10A-AMPKAR2-RAMPKAR2 | This report | N/A |
| MCF10A-AMPKAR2-RAMPKAR2-T/A | This report | N/A |
| MCF10A-RAMPKAR2-HYlight-mOr-nls | This report | N/A |
| MCF10A-RAMPKAR2-PercevalHR-nes | This report | N/A |
| MEF-WT |  |  |
| MEF-AMPKα1α2-double-KO | | |
| MEF-WT-PercevalHR-RAMPKAR2 | This report | N/A |
| MEF-AMPKα-dKO-PercevalHR-nls-RAMPKAR2 | This report | N/A |
| MEF-WT-RAMPKAR2-HYlight-mOr-nls | This report | N/A |
| MEF-AMPKα-dKO-RAMPKAR2-HYlight-mOr-nls | This report | N/A |
| **Recombinant DNA** |  |  |
| pPBJ-AMPKAR2-puro | Hung et al., 2017 | N/A |
| CMV-HYlight | Koberstein et al., 2022 | Addgene# 193447 |
| pPBJ-pHred-Perceval2-nls-puro |  |  |
| pBJ-Perceval2-nes |  |  |
| pLenti-RAMPKAR2-HYlight-mOr-nls | This report |  |
| pPBJ-RAMPKAR2-nes-neo | This report | Addgene# 212103 |
| pPBJ-RAMPKAR2-TA-nes-neo | This report | Addgene# 212104 |
| **Software and Algorithms** |  |  |
| NIS-Elements AR ver. 4.20 | Nikon | SCR_014329 |
| MATLAB | Mathworks | SCR_001622 |
| Bio-Formats | OME | SCR_000450 |
| **Other** |  |  |
| Glass bottom plates, #1.5H high-performance cover glass | Cellvis | P96-1.5H-N, P06-1.5H-N |

### Resource Availability

Further information and requests for resources and reagents should be directed to the lead contact, John Albeck, who will fulfill them.

### Materials and Availability

RAMPKAR2 and RAMPKAR2-T/A plasmids are available from Addgene (plasmid numbers: 212103 and 212104, respectively). Otherwise, cell lines and plasmids can be provided upon request, which must be made to John Albeck.

### Data and Code Availability

Image data collected via NIS elements can be made available upon request but could not be uploaded due to their large size (50-500 GB per ND2 file).

All data processing was performed in MATLAB using previously described methods ^44–47^. All code written to filter live-cell data, pool technical/experimental replicates, perform statistical analysis, and produce figures is available on GitHub at: https://github.com/Albeck-Lab/RAMPKAR

### Reporter construction

pPBJ-RAMPKAR2-neo was constructed by replacing the mTurq2 and YPet fluorophores in the previously described AMPKAR2 biosensor^7^, with miRFP670nano3^31^ and miRFP720^48^, respectively. The negative control version, pPBJ-RAMPKAR2-T/A-neo, was constructed by substituting the threonine residue in the AMPK motif substrate with an alanine residue, thus preventing phosphorylation and activation of the reporter.

### Reporter cell line generation

Stable cell lines were created by transfecting MCF10A (clone 5e) or MEF (either WT or AMPKαKO) with pBJ-PercevalHR-puro and/or pBJ-RAMPKAR2-neo combined with the PiggyBac retrotransposase system in FuGene4K. Cells were selected with neomycin (250 ug/ml for 7 days) and puromycin (3 ug/ml for 3 days), then cloned by limiting dilution. MCF10A and both MEF blank cell lines were also transduced using lentivirus bearing the RAMPKAR2-nes-HYlight-nes-mOrange-nls construct and cloned through limiting dilution. Two MEF AMPKαKO clones were then recombined to rule out accidental selection of a broken reporter.

### Cell Culture and Media

Regular cell culture of human mammary epithelial cells, MCF10A clone 5E^49^, was performed as previously described^50^. MCF10A cells were grown in supplemented DMEM/F12 (see Media table). Low passage stocks (< passage 3) from the parental MCF10A-5E clone were used to create all RAMPKAR2-containing cell lines, and previously published ‘MCF10A-AMPKAR2’ cell lines were used for the AMPKAR-RAMPKAR2/RAMPKAR2-T/A experiments. MEF cell lines (wild type or AMPKα1α2 double knockouts) were cultivated in DMEM high glucose GlutaMAX (see Media table). Cells were discarded after reaching a maximum of 20 passages from the initial parental MEF cultures. All cells were routinely passaged before reaching 85% confluency.

Live-cell imaging was performed using custom media (UC Davis Veterinary Medicine Biological Media Service) depleted of phenol red, riboflavin, and folic acid to eliminate background fluorescence. For MCF10A cells, we used ‘Imaging base-DMEM/F12’, which is further depleted of glucose and glutamine. All MCF10A experiments were performed in ‘MCF10A Imaging Medium’ (see *Media Composition*), which was supplemented with insulin and/or EGF when pre-treatments were required. For MEF cells, experiments utilized custom DMEM lacking the aforementioned background-contributing compounds (see *MEF Imaging Media Table*).

### Media Composition

**Table 1.** MCF10A Growth Medium.

| Reagent | Supplier | Item # | Stock Concentrations | Final Concentrations |
| --- | --- | --- | --- | --- |
| DMEM/F12 | Gibco | 11320033 | 1X | 1X |
| Horse Serum | Gibco | 26050088 | 100 % v/v | 5% v/v |
| Cholera Toxin | Sigma | C8052 | 1 mg/mL | 100 ng/mL |
| EGF | Peprotech | AF-100-15 | 100 µg/mL | 5 ng/mL |
| Hydrocortisone | Sigma | H0888 | 1 mg/mL | 0.5 µg/mL |
| Insulin | Sigma | I9278 | 10 mg/mL | 10 µg/mL |
| Biliverdin* | Sigma | 30891 | 32 mM | 25 µM |

\*Only supplemented into growth medium 12-24 hours prior to use for an imaging experiment.

**Table 2.** MEF Growth Medium.

| Reagent | Supplier | Item # | Stock Concentrations | Final Concentrations |
| --- | --- | --- | --- | --- |
| DMEM GlutaMAX high glucose | Gibco | 10564011 | 1X | 1X |
| Foetal Bovine Serum | GeminiBio | 100106500 | 100 % v/v | 9% v/v |
| Biliverdin* | Sigma | 30891 | 32 mM | 25 µM |

**Table 3.** MCF10A Imaging Medium.

| Reagent | Supplier | Item # | Stock Concentrations | Final Concentrations |
| --- | --- | --- | --- | --- |
| Imaging base-DMEM/F12* | UC Davis<br>Veterinary<br>Medicine<br>Biological<br>Media<br>Services |  | 1X | 1X |
| Bovine Serum Albumin | Sigma | A7906 | 1X | 0.2% w/v |
| Glucose | Fisher | A16828 | 1.74 M | 17 mM |
| Pyruvate | Gibco | 11360070 | 100 mM | 1 mM |
| Hydrocortisone | Sigma | H0888 | 1 mg/mL (2.76mM) | 0.5 ug/mL (1.38 µM) |
| Cholera Toxin | Sigma | C8052 | 1 mg/mL | 100 ng/mL |
| **<u>Optionally added:</u>** |  |  |  |  |
| Insulin | Sigma | I9278 | 10 mg/mL | 10 µg/mL |
| EGF | Peprotech | AF-100-15 | 100 µg/mL | 20 ng/mL |

\* Custom F12/DMEM similar to GIBCO 11320033 lacking glucose, glutamine, pyruvate, folic acid, riboflavin, and phenol red.

**Table 4.** MEF Imaging Medium.

| Reagent | Supplier | Item # | Stock Concentrations | Final Concentrations |
| --- | --- | --- | --- | --- |
| Imaging base - DMEM* | UC Davis<br>Veterinary<br>Medicine<br>Biological<br>Media<br>Services |  | 1X | 1X |
| Bovine Serum Albumin | Sigma | A7906 | 1X | 0.2% w/v |

\* Custom DMEM similar to GIBCO 1196511 lacking glucose, glutamine, pyruvate, folic acid, riboflavin, and phenol red.

**Table 5.** MCF10A Resuspension Medium.

| Reagent | Supplier | Item # | Stock Concentrations | Final Concentrations |
| --- | --- | --- | --- | --- |
| DMEM/F12 | Gibco | 11320033 | 1X | 1X |
| Horse Serum | Gibco | 26050088 | 1X | 10% v/v |

#### Live-cell fluorescence microscopy

Live-cell imaging experiments were conducted as described previously^47,51^. MCF10A and MEF cells were plated onto 96-well imaging plates more than 48h before imaging. Cells were treated with 25 μM biliverdin 12h before imaging and transferred into imaging media around 4h before the experiment started. Cells were imaged on a Nikon Ti-E inverted microscope with a stage-top incubator (37 °C, 5% CO2). Images were captured every six minutes using a Teledyne Photometrics Kinetix scMOS camera, a 20x/0.75 NA objective, and a Lumencor SPECTRA III light engine. Experiments used the following fluorescence filter sets: CFP (#49,001, Chroma), YFP (#49,003, Chroma), RFP670 (custom T647lpxr – ET667/30 m, Chroma, with the 635/22 nm SPECTRA III excitation band) and FRET720 (custom T647lpxr – FF01-730/39–32, Chroma and Semrock, with the 635/22 nm SPECTRA III excitation band). AMPKAR2 was measured using CFP and YFP filters, and RAMPKAR2 used the miRFP670 and FRET720 filters.

#### Signal cross-correlation

To quantify intracellular synchronization, we computed the normalized cross-correlation between paired single-cell fluorescence trajectories of RAMPKAR2 and either PercevalHR or HYlight. Analysis was restricted to defined temporal windows, such as the three-hour post-treatment interval, and explicitly excluded vehicle or off-target control conditions. After omitting missing values, traces were mean-centered to eliminate baseline offsets. We utilized the MATLAB `xcorr` function to calculate the normalized cross-correlation sequence for each cell, extracting the maximum absolute correlation coefficient and its corresponding temporal lag in minutes. To generate a null distribution for statistical comparison, we computed a paired scrambled control by randomly shuffling the timepoints of the first sensor trace using a reproducible seed before mean-centering and correlating it against the intact second sensor.

#### Pulse analysis, peak detection, and feature extraction

To comprehensively quantify biosensor responses, we extracted continuous trace kinetics and discrete pulse properties for each tracked cell. We defined specific analysis windows comprising a pre-treatment baseline period and a strictly bound post-treatment interval. From the continuous traces, we calculated the baseline mean, baseline signal variance, absolute treatment-induced signal shift, maximum post-treatment intensity, and the initial kinetic slope of the response. To identify individual oscillatory events, the MATLAB `findpeaks` function was applied to each cell’s biosensor trace to locate local maxima. Based on these identified peaks, the pipeline extracted supplementary peak-specific metrics. These included the total number of detected peaks, mean pulse duration, mean pulse amplitude, and the temporal delay to the first pulse following treatment.

#### Automated single-cell oscillation classification

Oscillatory dynamics in HYlight and Perceval biosensor channels were classified independently using a custom automated algorithm. We first excluded temporal traces shorter than 15 hours. For each experimental replicate, the classifier was manually trained on approximately 500 cell traces. Following training, we applied a prominence-based peak detection method to the full dataset, where minimum peak distance constrained the biologically plausible period and prominence thresholds scaled to cell-specific signal variability. The algorithm computed three kinetic metrics per trace: the spectral power fraction within the defined period window, the coefficient of variation of inter-peak intervals, and overall signal amplitude. These features generated a continuous oscillatory score ranging from 0 to 1. Based on this continuous score and the persistence of detected peaks, cells were assigned to one of three classes: non-oscillatory (no significant peaks), medium oscillatory (intermittent peaks), or strong oscillatory (sustained periodic peaks). For each experimental replicate, the classifier was manually trained until it reached >70% accuracy.

#### Similarity-ordered row heatmaps

Raw single-cell trajectories were row-scaled using Z-score standardization. Each trace was mean-centered and divided by its standard deviation and computationally clipped at a maximum threshold of three standard deviations to mitigate the visual impact of extreme outliers. To group similar temporal patterns, these scaled trajectories were hierarchically clustered within their assigned oscillation class. This clustering utilized a correlation-based distance metric and average linkage, followed by optimal leaf ordering to ensure adjacent rows exhibited maximal dynamic similarity.

### Biosensor quantification

Biosensor signals were interpreted from image intensity via models reflecting the molecular function of each sensor and the spectral modulation of the microscope light path. To enable consistent measurements, the light path was characterized by manufacturer-provided light source and filter spectra combined with on-site collection of calibration data of light intensity through the relevant filters. In each case, the image intensities were used to estimate the fraction of reporter molecules that are activated or bound to their target.

AMPKAR2 and RAMPKAR are both FRET reporters, so their signals were estimated based on a previous model^46^ (originally established for ERK kinase activity reporter, EKAR). RAMPKAR was modeled on the same principles, but with different properties reflecting the different fluorophores used, and using a different spectral measurement scheme (FRET/Donor, rather than Donor/Acceptor). This model has been recently adapted for red-shifted FRET reporters for ERK^29^, which employ the same fluorophores as RAMPKAR.

In contrast, PercevalHR and HYlight are both spectral sensors, based on fusions with circularly permuted GFP, such that binding of the target molecule biases the spectral properties of the fluorophore. The signal strength of these reporters was modeled via a mixture model representing the two molecular states: ADP- vs. ATP-bound for PercevalHR, and free vs. FBP-bound for HYlight. For clarity, the derivation of this model is presented below, starting from the fluorescent imaging model of Gillies et al.^46^, appendix, Eqn. 1.

With two species of fluorophore in the imaging control volume (i.e. per pixel), the image intensity for a given excitation/emission scheme is given by eqn. 1, defining the following terms

*I* : Imaging intensity

*P* : Excitation power gain

*t* : Exposure time

*C*_*F*_, *C*_*B*_ : Concentrations of free and bound sensor, respectively

*X*_*F*_(λ), *X*_*B*_(λ) : Excitation spectra, depends on wavelength (λ), light source spectrum, filter spectra, and fluorophore absorbance spectrum

*M*_*F*_(λ), *M*_*B*_(λ) : Emission spectra, including filters and camera quantum efficiency

*Q*_*F*_, *Q*_*B*_ : Quantum yields

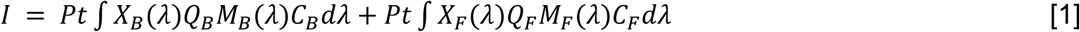

Since only two states of the sensor exist in the model, free and bound, the concentrations may be reframed in terms of the fractions of sensor free and bound, via the definitions:

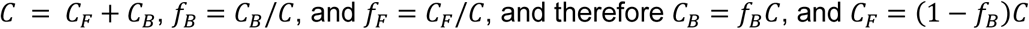

Making these substitutions and evaluating the integrals over the spectra of the light source, filters, fluorophores and camera yields eqn. 2.

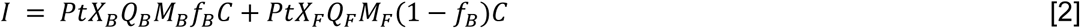

For convenience, we define summary terms that depend on the excitation scheme and which may be computed based on calibration data:

*L*_*F*_ = *X*_*F*_*Q*_*F*_*M*_*F*_, and *L*_*B*_ = *X*_*B*_*Q*_*B*_*M*_*B*_ when excited with the “long wavelength” channel, and *S*_*F*_ = *X*_*F*_*Q*_*F*_*M*_*F*_, *S*_*B*_ = *X*_*B*_*Q*_*B*_*M*_*B*_, when excited with the “short wavelength” channel.

Substituting with these summary terms, we can concisely consider the ratio of the “long” channel over the “short”, eqn. 3. In this ratio, the common terms of power, time and total sensor concentration all cancel.

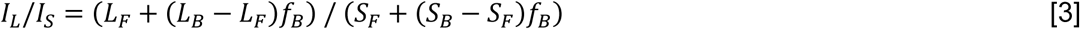

Solving for the fraction of sensor bound, *fB*, yields the final estimate of the sensor signal, eqn. 4

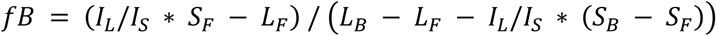

## Figure Legends

**Supplemental figure 1:**
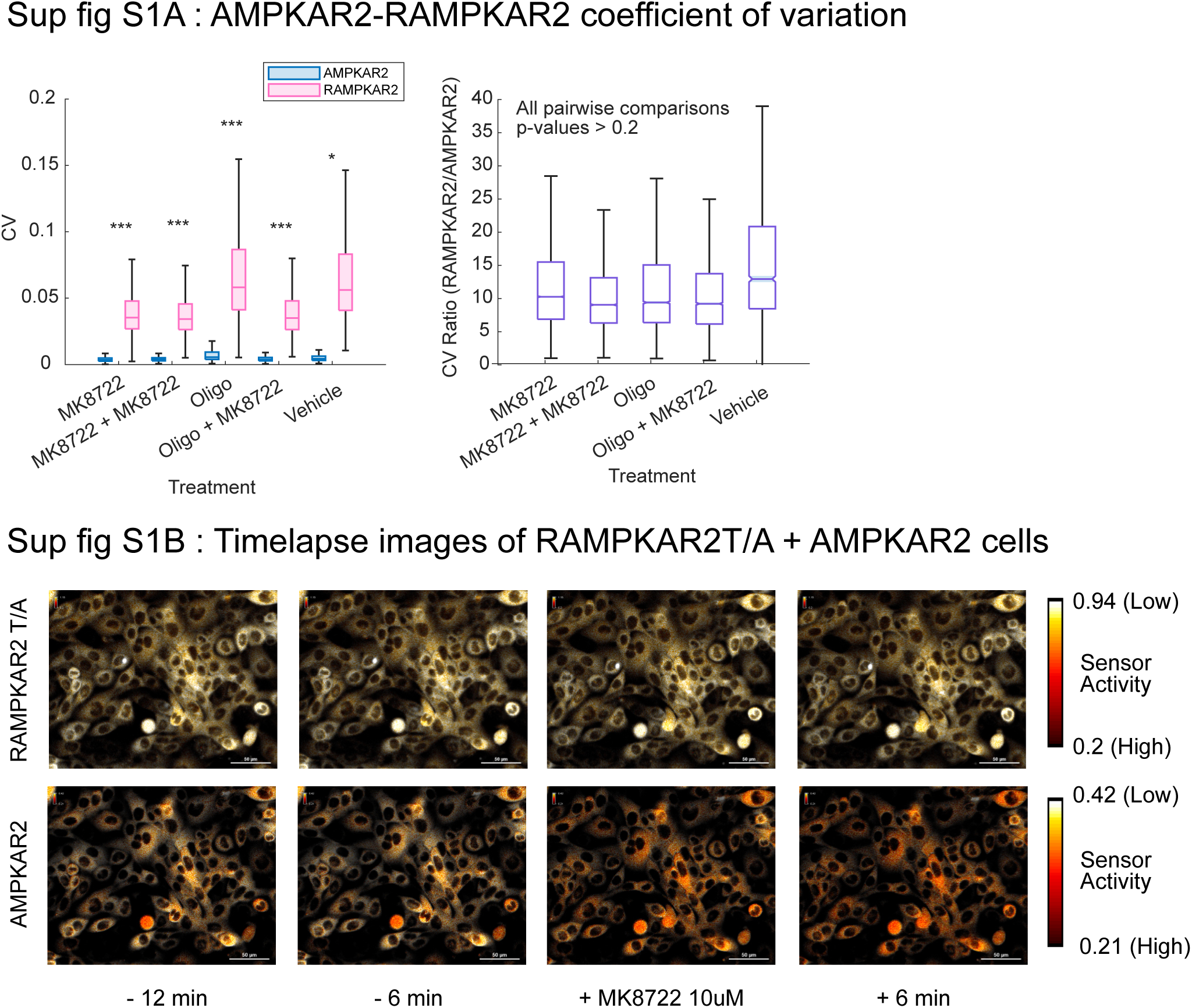
**(A)** Coefficient of variation (CV) between AMPKAR2 and RAMPKAR2 signals in stable time windows, per treatment (left panel) and ratio of the coefficient of variation (CV) (right panel). RAMPKAR2 exhibits significantly higher CV than AMPKAR2 across all treatment conditions (paired t-test, cell-level with > 1800 cells per condition, * = p<0.05 and *** = p<0.0001). CV ratios (RAMPKAR2/AMPKAR2) are stable across all treatment conditions with no significant pairwise differences (two-sample pairwise t-test, replicate-level with n = 3, all p-values>0.2). **(B)** Example time-lapse images of the RAMPKAR-T/A and AMPKAR2 (top and bottom rows) signals, in the same MCF-10A cells, before and after treatment with the AMPK activator MK8722. Top row: Images of the RAMPKAR-T/A FRET ratio over time, with colour bar indicating signal. Bottom row: AMPKAR2 images of the same cells as the top row, with associated colour bar indicating signal. For RAMPKAR-T/A and AMPKAR2, relative AMPK activity is represented in pseudo-colouring, with low relative AMPK activity in white and high AMPK activity in red. Scale bar is 50 μm.

**Supplemental figure 2:**
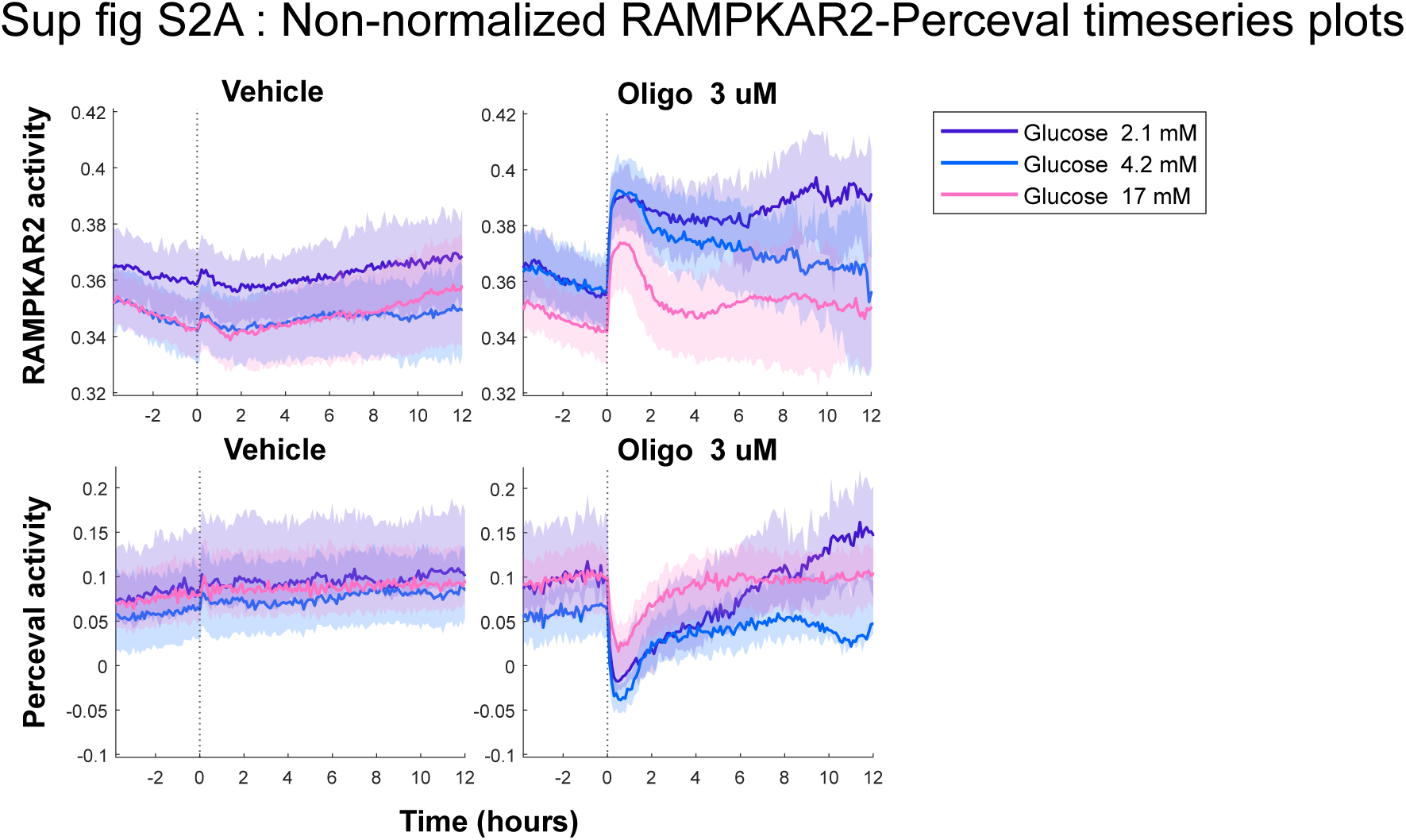
**(A)** Non-baseline normalized timeseries from RAMPKAR2-PercevalHR MCF-10A cells. Cells stably expressing the biosensor combination were cultured in different glucose concentrations and treated with oligomycin 3 uM. These plots represent the same data shown in Figure 2, but without normalization to the baseline signal. Due to low expression of PercevalHR within these cells, variation in baseline signal was subject to variation based on minor differences in the regions used for background subtraction, resulting in potentially artifactual differences in the signal amplitude following oligomycin treatment. Rather than attempt to optimize background subtraction with potentially subjective choices in background subtraction regions, we provide here the non-normalized data alongside the main figure data that include baseline normalization to better emphasize the differences in oligomycin-stimulated changes in signal.

**Supplemental figure 3:**
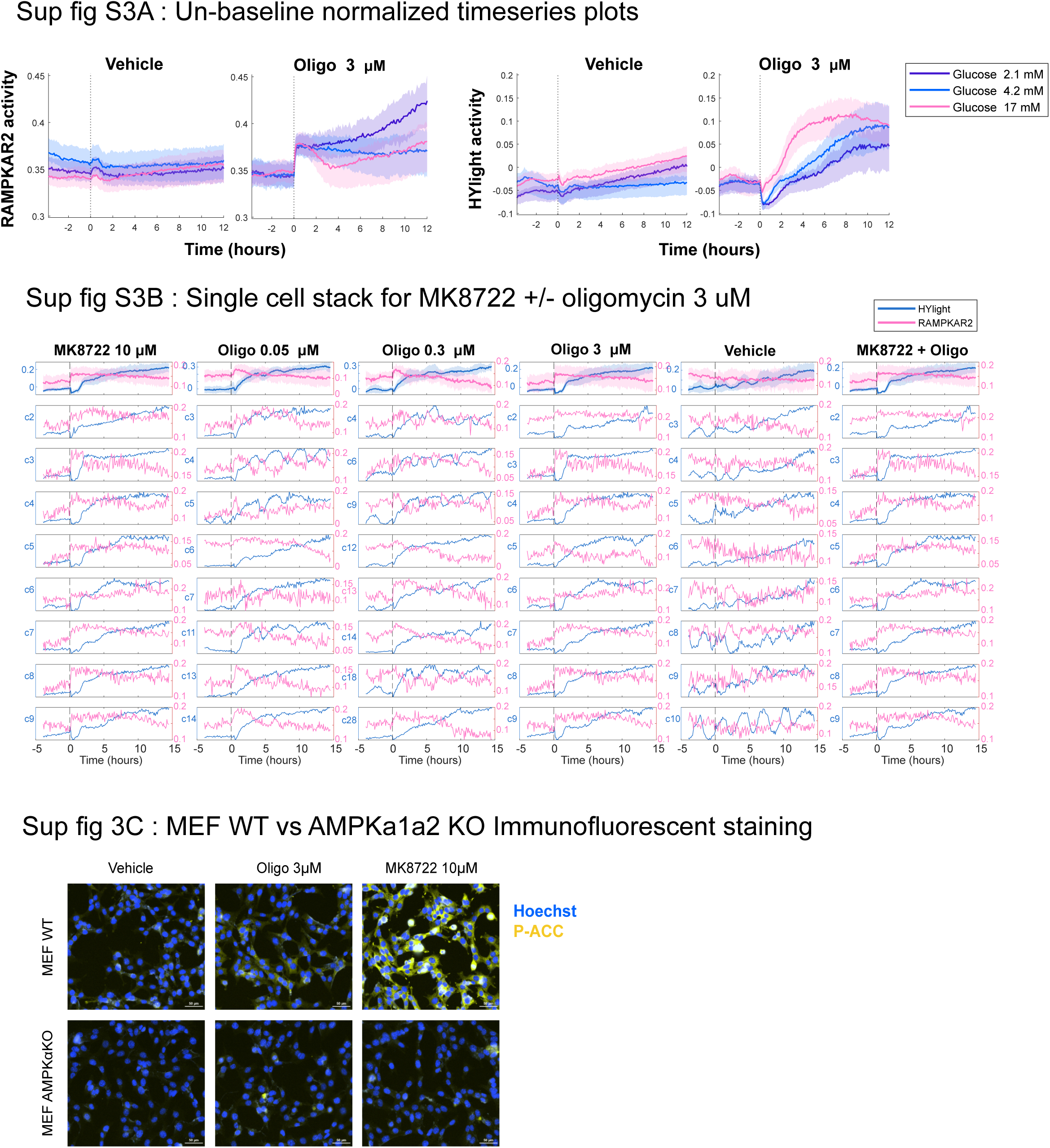
**(A)** Non-baseline normalized timeseries from RAMPKAR2-HYlight MCF-10A cells. These plots represent the same data shown in Figure 3A and 3B. While biosensor expression was not low and baseline subtraction was less of an issue for this biosensor cell line than for RAMPKAR2-PercevalHR (Figure 2 and Figure S2), we present both the normalized and non-normalized data for consistency of presentation across the cell lines. **(B)** Single-cell traces from one RAMPKAR2-HYlight MCF-10A experimental replicate. MK8722 +/- oligomycin response pattern is consistent cell-by-cell as opposed to response to oligomycin treatments. Left y axis indicates the number of the cell trace shown as well as HYlight sensor units in the first row. Right y axis indicates RAMPKAR2 sensor activity. **(C)** MEF WT and AMPKa1a2 dKO cultured in DMEM 17mM glucose for 3 hours and treated with either oligomycin 3 uM or MK8722 10 μM. Cells were fixed and stained 2 hours after treatment with Hoechst (blue, nuclear) or phospho-ACC (yellow, cytoplasmic). Image area was chosen within the center of the image and applied for all subsequent cropped image. Scale bar is 50 μm. Images shown are from one of two technical replicates within one of two biological replicates (n = 2, > 400 cells per condition).

## Author Contributions

Project conception: NLD, JGA. Reagent development: NLD, MH, MBC, JH, CA. Data acquisition: NLD, MH, MBC. Data analysis: MH, NLD, MP. Data interpretation: JGA, NLD, MH, MP. Figure making: MH, NLD, MP. Manuscript writing: JGA, NLD. Manuscript editing: JGA, MH, MP, NLD

