## Supplemental figure 1 for "A Far-Red FRET biosensor for AMPK enables multiplexed imaging of single-cell bioenergetic homeostasis"

### Sup fig S1A : AMPKAR2-RAMPKAR2 coefficient of variation

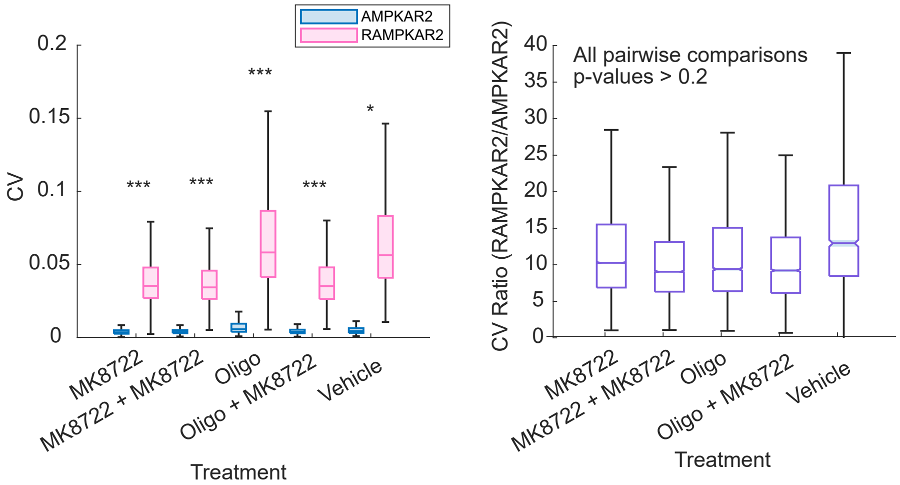


### Sup fig S1B : Timelapse images of RAMPKAR2T/A + AMPKAR2 cells

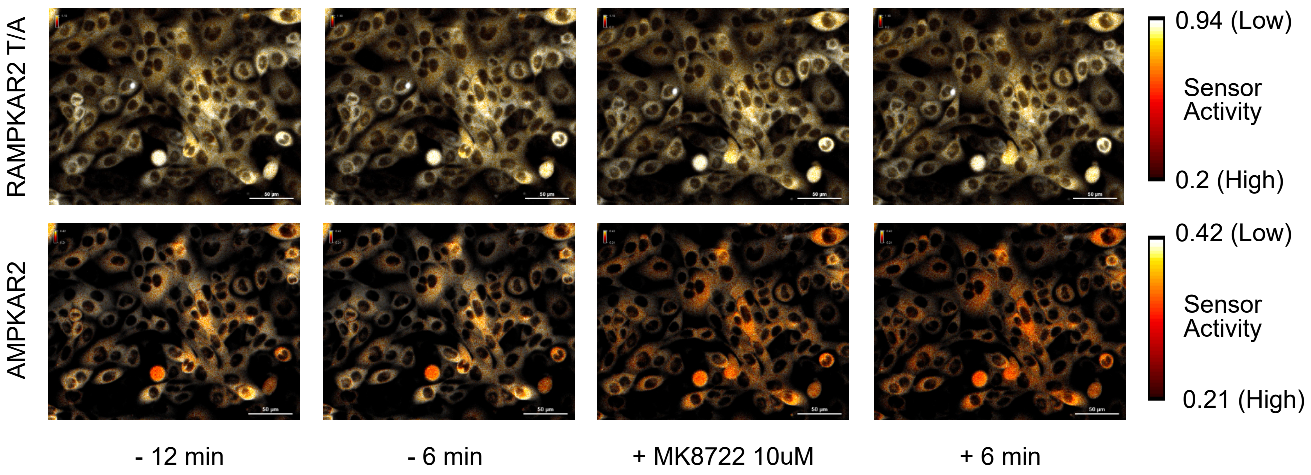


**Supplemental figure 1**: **(A)** Coefficient of variation (CV) between AMPKAR2 and RAMPKAR2 signals in stable time windows, per treatment (left panel) and ratio of the coefficient of variation (CV) (right panel). RAMPKAR2 exhibits significantly higher CV than AMPKAR2 across all treatment conditions (paired t-test, cell-level with > 1800 cells per condition, * = p<0.05 and *** = p<0.0001). CV ratios (RAMPKAR2/AMPKAR2) are stable across all treatment conditions with no significant pairwise differences (two-sample pairwise t-test, replicate-level with n = 3, all p-values>0.2). **(B)** Example time-lapse images of the RAMPKAR-T/A and AMPKAR2 (top and bottom rows) signals, in the same MCF-10A cells, before and after treatment with the AMPK activator MK8722. Top row: Images of the RAMPKAR-T/A FRET ratio over time, with colour bar indicating signal. Bottom row: AMPKAR2 images of the same cells as the top row, with associated colour bar indicating signal. For RAMPKAR-T/A and AMPKAR2, relative AMPK activity is represented in pseudo-colouring, with low relative AMPK activity in white and high AMPK activity in red. Scale bar is 50 μm.
