## Supplemental figure 2 for "A Far-Red FRET biosensor for AMPK enables multiplexed imaging of single-cell bioenergetic homeostasis"

### Sup fig S2A : Non-normalized RAMPKAR2-Perceval timeseries plots

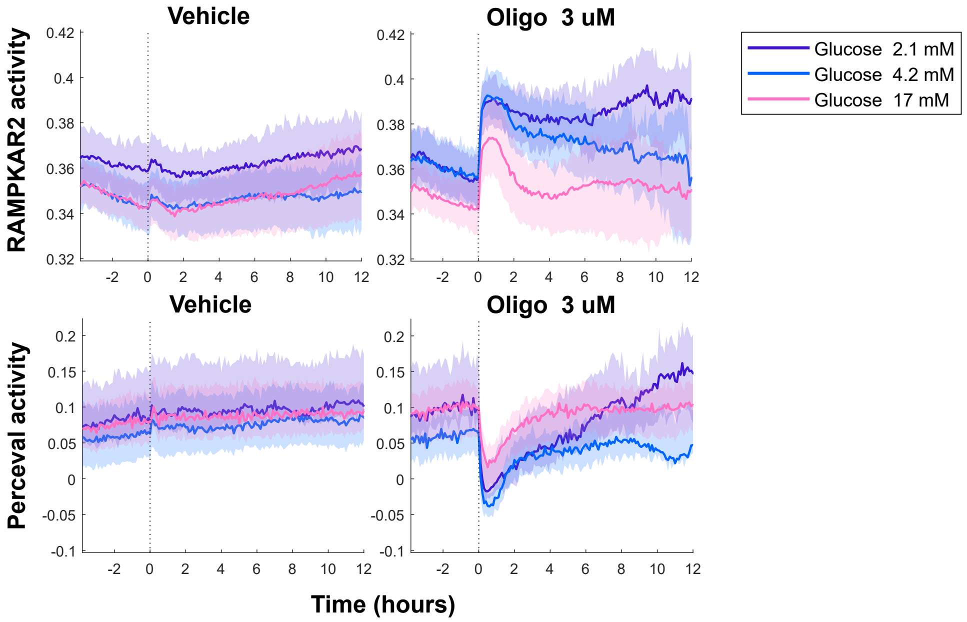


**Supplemental figure 2: (A)** Non-baseline normalized timeseries from RAMPKAR2-PercevalHR MCF-10A cells. Cells stably expressing the biosensor combination were cultured in different glucose concentrations and treated with oligomycin 3 µM. These plots represent the same data shown in Figure 2, but without normalization to the baseline signal. Due to low expression of PercevalHR within these cells, variation in baseline signal was subject to variation based on minor differences in the regions used for background subtraction, resulting in potentially artifactual differences in the signal amplitude following oligomycin treatment. Rather than attempt to optimize background subtraction with potentially subjective choices in background subtraction regions, we provide here the non-normalized data alongside the main figure data that include baseline normalization to better emphasize the differences in oligomycin-stimulated changes in signal.
