## Supplemental figure 3 for "A Far-Red FRET biosensor for AMPK enables multiplexed imaging of single-cell bioenergetic homeostasis"

### Sup fig S3A : Un-baseline normalized timeseries plots

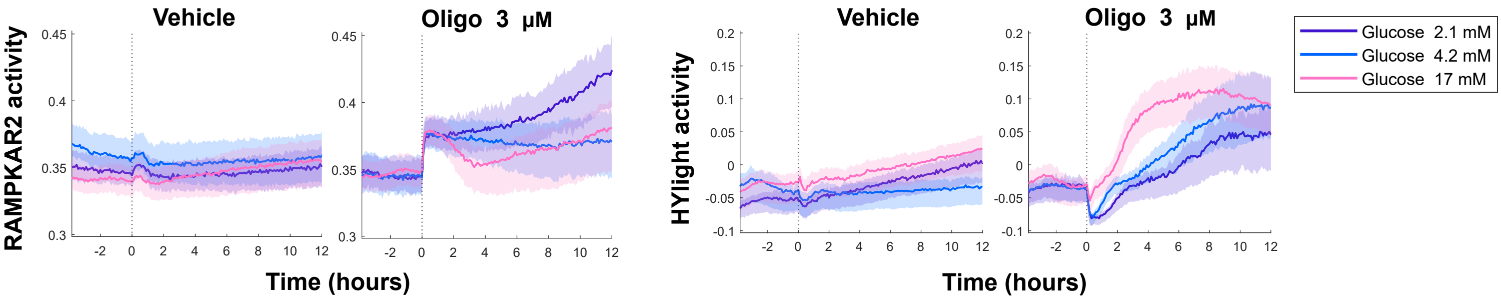


### Sup fig S3B : Single cell stack for MK8722 +/- oligomycin 3 uM

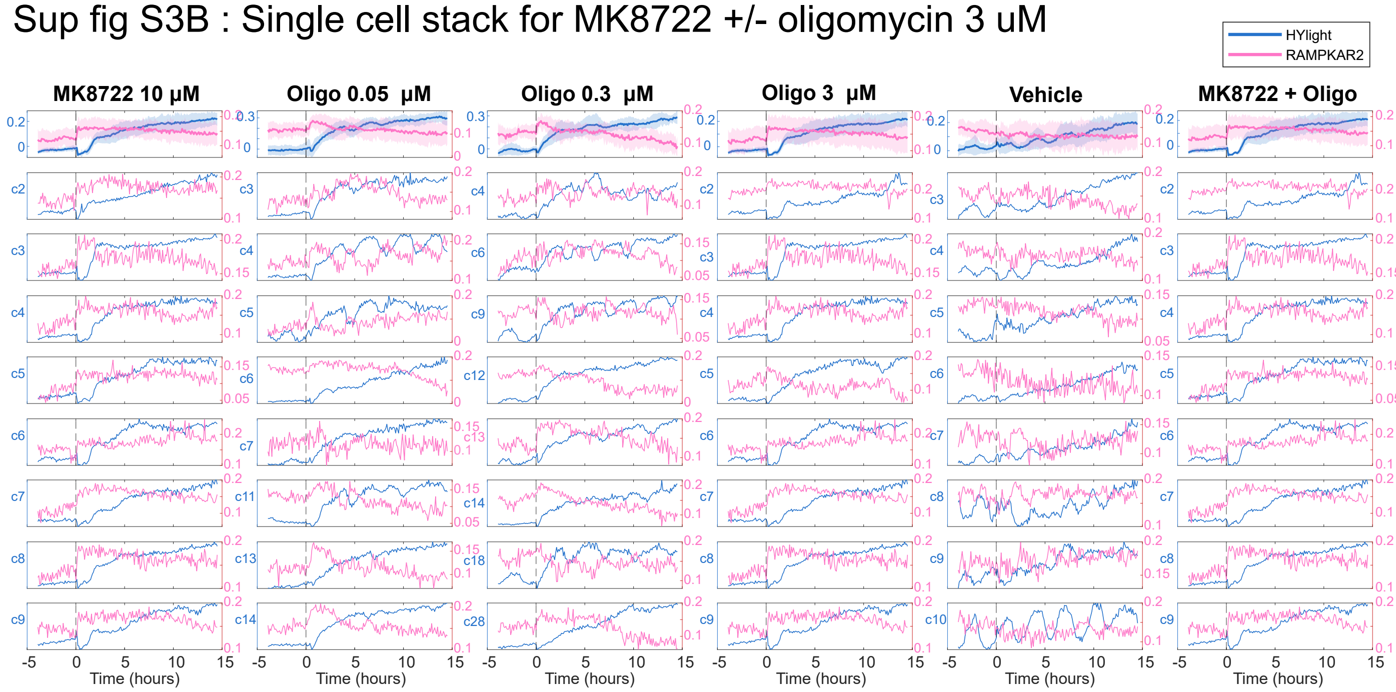


### Sup fig 3C : MEF WT vs AMPKa1a2 KO Immunofluorescent staining

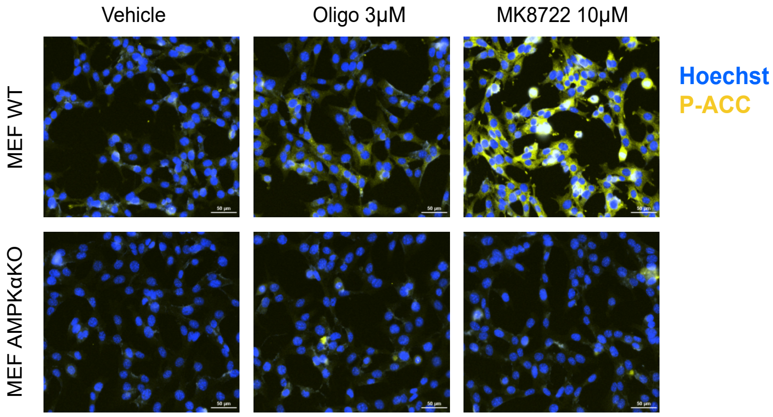


**Supplemental figure 3: (A)** Non-baseline normalized timeseries from RAMPKAR2-HYlight MCF-10A cells. These plots represent the same data shown in Figure 3A and 3B. While biosensor expression was not low and baseline subtraction was less of an issue for this biosensor cell line than for RAMPKAR2-PercevalHR (Figure 2 and Figure S2), we present both the normalized and non-normalized data for consistency of presentation across the cell lines. **(B)** Single-cell traces from one RAMPKAR2-HYlight MCF-10A experimental replicate. MK8722 +/- oligomycin response pattern is consistent cell-by-cell as opposed to response to oligomycin treatments. Left y axis indicates the number of the cell trace shown as well as HYlight sensor units in the first row. Right y axis indicates RAMPKAR2 sensor activity. (C) MEF WT and AMPKa1a2 dKO cultured in DMEM 17mM glucose for 3 hours and treated with either oligomycin 3 uM or MK8722 10 μM. Cells were fixed and stained 2 hours after treatment with Hoechst (blue, nuclear) or phospho-ACC (yellow, cytoplasmic). Image area was chosen within the center of the image and applied for all subsequent cropped image. Scale bar is 50 μm. Images shown are from one of two technical replicates within one of two biological replicates (n = 2, > 400 cells per condition).
